# Relating anthropometric indicators to brain structure in 2-month-old Bangladeshi infants growing up in poverty: a pilot study

**DOI:** 10.1101/655068

**Authors:** Ted Turesky, Wanze Xie, Swapna Kumar, Danielle D. Sliva, Borjan Gagoski, Jennifer Vaughn, Lilla Zöllei, Rashidul Haque, Shahria Hafiz Kakon, Nazrul Islam, William A. Petri, Charles A. Nelson, Nadine Gaab

**Affiliations:** Laboratories of Cognitive Neuroscience, Division of Developmental Medicine, Department of Medicine, Boston Children’s Hospital, Boston, Massachusetts, U.S.; Harvard Medical School, Boston, Massachusetts, U.S.; Department of Neuroscience, Brown University, Providence, Rhode Island, U.S.; Department of Radiology, Harvard Medical School, Boston, Massachusetts, U.S.; Fetal Neonatal Neuroimaging and Developmental Science Center, Boston Children’s Hospital, Boston, Massachusetts, U.S.; A.A. Martinos Center for Biomedical Imaging, Massachusetts General Hospital, Boston, Massachusetts, U.S.; The International Centre for Diarrhoeal Disease Research, Dhaka, Bangladesh; National Institute of Neuroscience and Hospital, Dhaka, Bangladesh; Division of Infectious Diseases and International Health, Department of Medicine, School of Medicine, University of Virginia, Charlottesville, Virginia, U.S.; Harvard Graduate School of Education, Cambridge, Massachusetts, U.S.

**Keywords:** infants, stunting, underweight, wasting, adversity, MRI

## Abstract

Anthropometric indicators, including stunting, underweight, and wasting, have previously been associated with poor neurocognitive outcomes. This link may exist because malnutrition and infection, which are known to affect height and weight, also impact brain structure according to animal models. However, a relationship between anthropometric indicators and brain structural measures has not been tested yet, perhaps because stunting, underweight, and wasting are uncommon in higher-resource settings. Further, with diminished anthropomorphic growth prevalent in low-resource settings, where biological and psychosocial hazards are most severe, one might expect additional links between measures of poverty, anthropometry, and brain structure. To begin to examine these relationships, we conducted an MRI study in 2-3-month-old infants growing up in the extremely impoverished urban setting of Dhaka, Bangladesh. The sample size was relatively small because the challenges of investigating infant brain structure in a low-resource setting needed to be realized and resolved before introducing a larger cohort. Initially, fifty-four infants underwent T_1_ sequences using 3T MRI, and structural images were segmented into gray and white matter maps, which were carefully evaluated for accurate tissue labeling by a pediatric neuroradiologist. Gray and white matter volumes from 29 infants (79 ± 10 days-of-age; F/M = 12/17), whose segmentations were of relatively high quality, were submitted to semi-partial correlation analyses with stunting, underweight, and wasting, which were measured using height-for-age (HAZ), weight-for-age (WAZ), and weight-for-height (WHZ) scores. Positive semi-partial correlations (after adjusting for chronological age and sex and correcting for multiple comparisons) were observed between white matter volume and HAZ and WAZ; however, WHZ was not correlated with any measure of brain volume. In examining the role of poverty, no associations were observed between income-to-needs or maternal education and brain volumetric measures, suggesting that risk factors previously linked with poverty were not associated with total brain tissue volume pre- or peri-natally in this sample. Overall, these results provide the first link between diminished anthropomorphic growth and white matter volume in infancy. Challenges of conducting a developmental neuroimaging study in a low-resource country are described.

## INTRODUCTION

Hundreds of millions of children worldwide fail to reach their full growth potential due to adverse circumstances in their environments (Bhutta et al., 2017; Black et al., 2017; Grantham-McGregor et al., 2007; John et al., 2017). To assess and monitor diminished growth, many nations have adopted the World Health Organization (WHO) anthropometric indicators: stunting, measured with height-for-age (HAZ); underweight, measured with weight-for-age (WAZ); and wasting, measured with weight-for-height (WHZ) (de Onis et al., 2012). Thus far, anthropometric indicators have been associated with poor neurocognitive outcomes (de Onis and Branca, 2016; Fuglestad et al., 2008) and increased risk of premature death (Black et al., 2013; de Onis and Branca, 2016; Hayashi et al., 2018), making them critical for identifying children at risk for later consequences (de Onis et al., 2012). Identification could be especially beneficial in areas such as Sub-Saharan Africa and Southern Asia, where the prevalence of diminished growth is especially high (Hayashi et al., 2018).

Nonetheless, these indicators may be limited in their utility to characterize other aspects of diminished growth, such as compromised brain development. While brain structure has not yet been examined in the context of anthropometric indicators, the link between stunting, underweight and wasting and neurocognitive outcomes (de Onis and Branca, 2016; Donowitz et al., 2018; Fuglestad et al., 2008) suggests that compromised brain structure and function may be associated with these anthropometric indicators.

Additional support for this assertion comes from a breadth of evidence showing that stunting and wasting are mainly caused by malnutrition and infection pre- and post-natally in the first two years of life (Black et al., 2013; de Onis and Branca, 2016; Schnee et al., 2018; Stewart et al., 2013), both of which are thought to also affect brain development (Hagberg et al., 2015; Keunen et al., 2015; Levitsky and Strupp, 1995; Steenwinckel et al., 2014). The relationship between these biological risk factors and anthropometric indicators is so well established that stunting, underweight, and wasting have been considered *forms* of malnutrition rather than only outcomes of it; e.g., “stunting is the most prevalent form of child malnutrition” (de Onis and Branca, 2016). Several different mechanisms could explain this association (Black et al., 2013; Stewart et al., 2013), including infection-induced inflammation (Jiang et al., 2014), and it has also been proposed that some of these also apply to brain structure (Jensen et al., 2017). For instance, deficiencies of nutrients (e.g., zinc) critical for basic biological processes (e.g., protein synthesis) would presumably impact both non-neural and neural processes. In addition, malnutrition and infection are likely to reciprocate harm (e.g., poorer appetite when sick, or, conversely, immune dysfunction due to lack of key nutrients; Rytter et al., 2014).

Several studies using animal models have directly examined the relationship between biological risk factors and brain development. At the cellular level, malnutrition has been associated with reductions in cortical cell size, density of cortical dendritic spines, myelin production, number of synapses, and number of glial cells (Levitsky and Strupp 1995; Keunen et al. 2015; for a review, please see Jensen et al., 2017). Meanwhile, infections have been consistently linked with white matter injury via inflammatory responses (Duggan et al., 2001; Hagberg et al., 2015; Hansen-Pupp et al., 2005). This latter link has been elucidated in a diffusion tensor imaging study by Favrais and colleagues (2011), who examined mice with systemic inflammation as caused by interleukin-1β (IL-1β), a pro-inflammatory cytokine. Mice with systemic inflammation exhibited reduced fractional anisotropy and radial diffusivity, measures of white matter microstructure, as well as reduced performance on long-term and working memory tasks, compared with control mice. Subsequent investigation of mechanism revealed that systemic inflammation interfered with normative maturation of oligodendrocytes (Favrais et al., 2011), which are required for axonal myelination in the central nervous system. Critically, these results provide mechanistic support for studies in humans, which have linked inflammation (Jensen et al., 2019a; Jiang et al., 2017, 2014) and deficiencies of nutrients such as phosphatidylcholine species (Moreau et al., 2019) in Bangladeshi infants from low- and middle-income families with later neurodevelopmental outcomes.

Based on these findings, we adapted a model in which biological hazards such as malnutrition and infection give rise to both anthropometric indicators of stunting, underweight, and wasting, and, in parallel, alterations in brain structure, which subsequently lead to poor neurocognitive outcomes (Jensen et al. 2017; Fig. 1). Testing a model such as this is difficult because current neuroimaging tools primarily utilized to measure brain structure and function (e.g., MRI, EEG) are mainly limited to middle- and high-resource research settings and populations, where severe malnutrition and infection are not common (Nelson, 2017).

**Figure 1.**
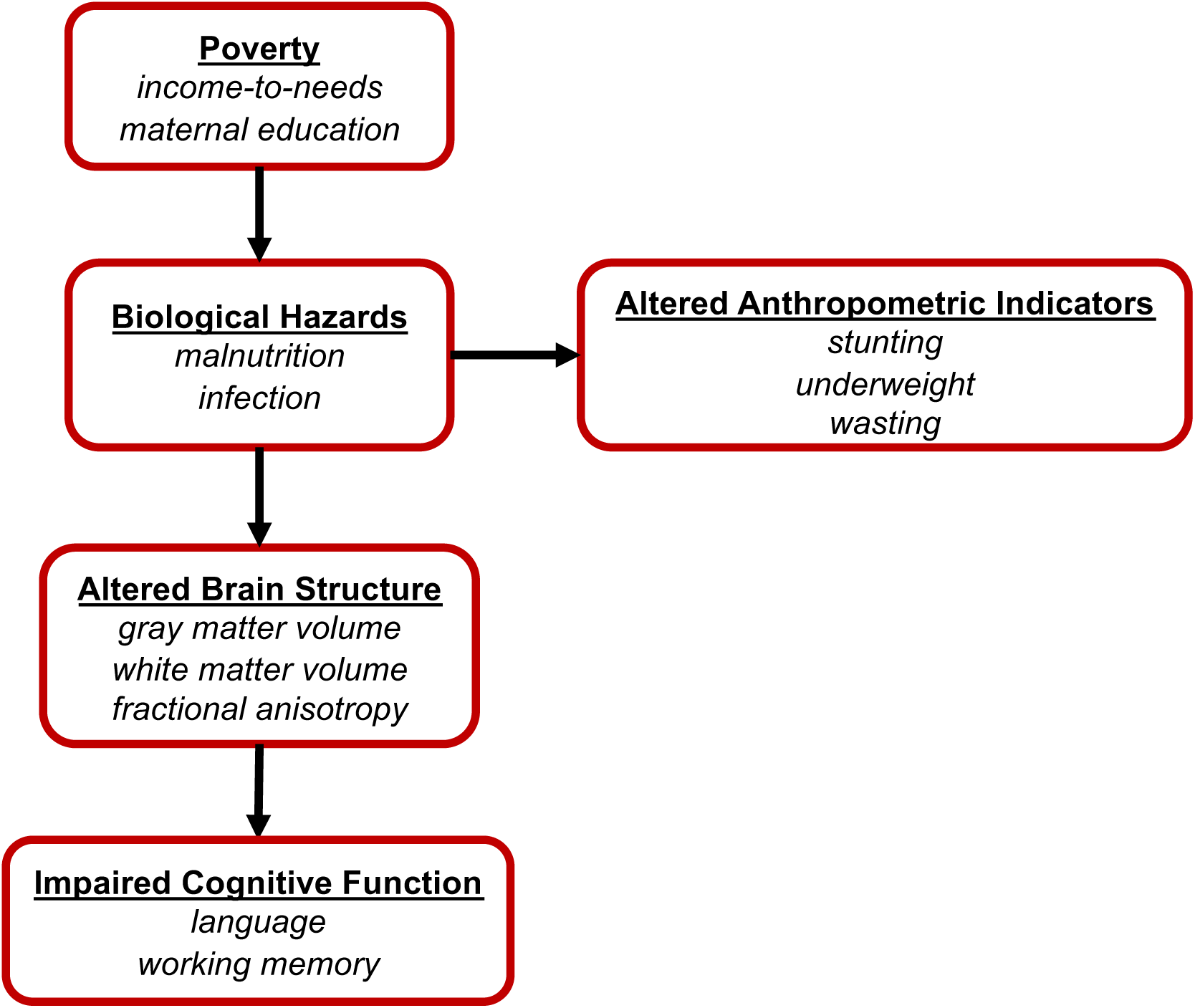
Hypothesized model illustrating the pathway through which poverty impairs cognitive function. Bolded, underlined text indicates risk factor categories, and italicized text indicates example risk factors within each category. This model is adapted from Jensen et al., 2017.

To resolve this, we set up a study in an extremely low-resource, urban area of Dhaka, Bangladesh (Fig. 2A), where biological hazards are severe. Measures were acquired at the age of 2 months, as infancy is the period of most rapid neural development (Gilmore et al., 2012), when children are most sensitive to environmental factors of nutrition, infection, and psychosocial caretaking (Black et al., 2013; de Onis and Branca, 2016; Jensen et al., 2017). As this was the first research study of infant brain structure in Bangladesh, several initial challenges needed to be overcome: e.g., an MRI facility needed to be identified and equipped for infant research scanning; compliance protocols needed to be aligned across collaborating institutions; and research staff required training on recruitment practices and infant imaging techniques. Further, it was not yet clear whether these challenges (for conducting an MRI study in a new location) would be intensified by the low-resource setting. As such, before introducing a larger cohort, this study was conducted on a small, pilot sample (for an examination of functional MRI in a low-resource setting, please see Turesky et al. 2019). This small sample size limited the components of the proposed model that could be tested.

**Figure 2.**
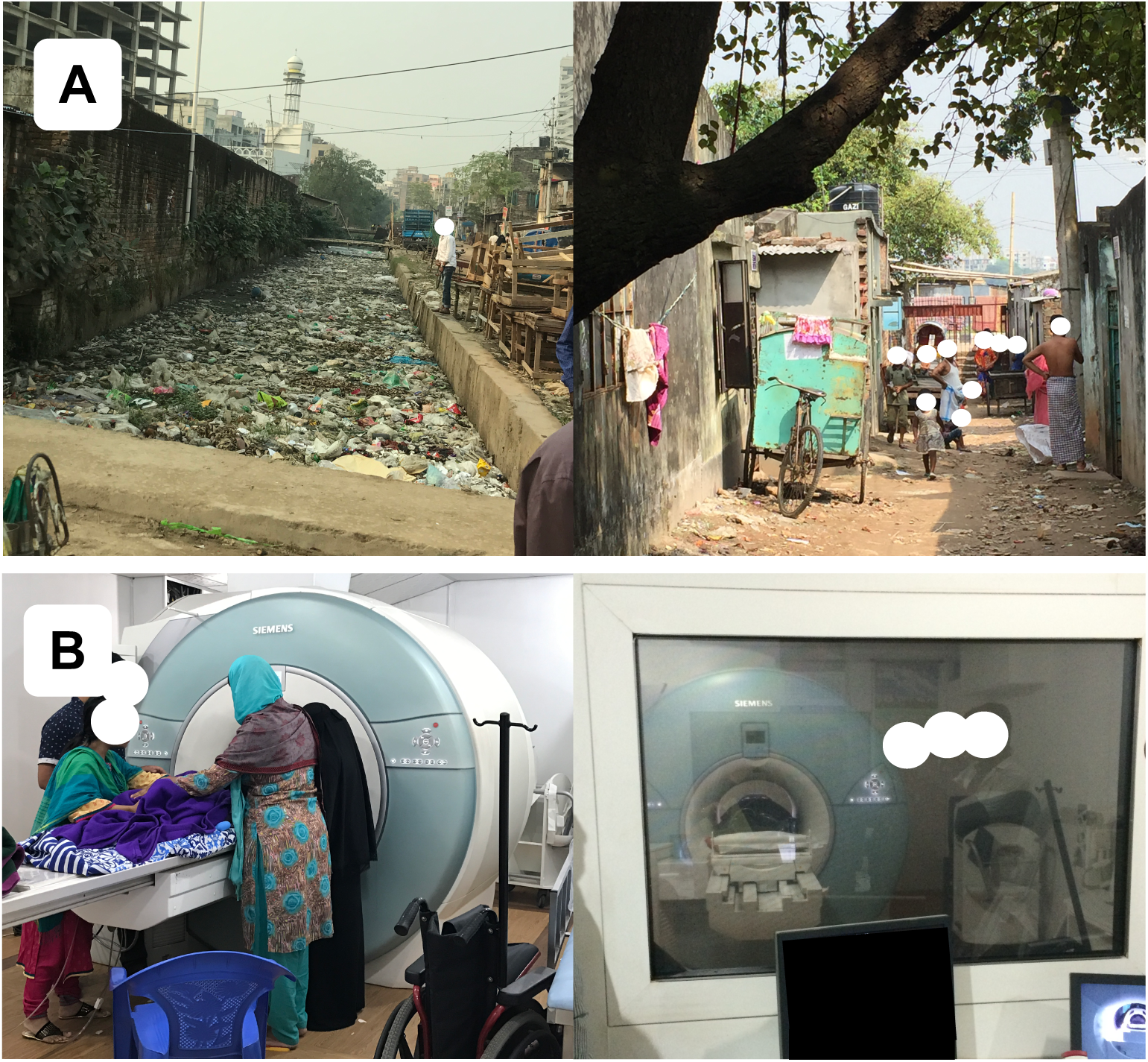
Study location. (A) Representative neighborhood in Dhaka, Bangladesh, where infants in this study live. (B) Facility at NINSH where infants undergo MRI scanning.

Anthropometric indicators of stunting, underweight, and wasting are reflective of malnutrition and infection (Black et al., 2013; de Onis and Branca, 2016; Stewart et al., 2013) and associated with poor neurocognitive outcomes (de Onis and Branca, 2016; Fuglestad et al., 2008). In addition, malnutrition and inflammation have been linked to alterations in brain properties (Duggan et al., 2001; Favrais et al., 2011; Hagberg et al., 2015; Hansen-Pupp et al., 2005; Keunen et al., 2015; Korzeniewski et al., 2014; Levitsky and Strupp, 1995; Malaeb et al., 2014; Yoon et al., 1997). Therefore, we hypothesized an association between anthropometry and brain structural measures. Furthermore, as these biological hazards affect measurements of the human body globally, one might expect that corresponding alterations in the brain would also occur globally, rather than to specific brain regions; therefore, structural measures of total gray and white matter volumes (G/WMV) were chosen for analyses.

Additionally, since stunting, underweight, and wasting are more likely to occur in low-resource settings beset by poverty, a secondary goal of this study was to determine whether brain structure measures were related to measures of poverty. The relationship between anthropometry and poverty seems to be age-dependent (Victora et al., 2010), such that anthropometric indicators might not reflect risk factors associated with poverty (e.g., malnutrition) before six months of age (Kerac et al., 2015). However, it is possible that measurements of brain structure are more sensitive compared with measurements of height and weight at two-three months-of-age and thus would be more likely to exhibit an association with measures of poverty, were one to exist. This would be consistent with a previous study in a partially overlapping sample of Bangladeshi infants, in which infants from families living in extreme poverty exhibited greater (i.e., less negative) intrinsic functional connectivity between the amygdala and precuneus compared with infants from relatively higher-income families (Turesky et al., 2019). In addition, previous studies of infants in the U.S. have reported a positive relationship between income-to-needs (a variable commonly used to measure poverty; or another income measure) and regional and total GMV (Betancourt et al., 2016; Hanson et al., 2013), but these links have not been directly tested in infants living in environments of extreme poverty, where biological and psychosocial hazards are most severe. Taken together, we hypothesized a negative association between poverty and brain volume. To test this, we examined the relationship between two measures of poverty, income-to-needs (i.e., household family income divided by number of household members) and years of maternal education, and G/WMV.

## METHODS

### Participants

The present study is a subset of the Bangladesh Early Adversity Neuroimaging (BEAN) study (Berens et al., n.d.; Jensen et al., 2019a,b; Moreau et al., 2019; Turesky et al., 2019), investigating effects of extreme biological and psychosocial adversity on early neurocognitive development in infants and children in Dhaka, Bangladesh (Fig. 2A). The broader study tracks 235 infants through early childhood using a variety of behavioral batteries and neuroimaging techniques, while the current study focused on the 54 infants who underwent structural MRI at two-months-of-age. Scanning was limited to a subset because the potential pitfalls associated with structural and functional neuroimaging in a low-resource setting needed to be elucidated before scanning a larger sample, as this was the first MRI study on infants in Bangladesh. Of 54 infants who attempted MRI, 53 completed the structural/MPRAGE sequence. One infant did not attempt it as they did not fall asleep within the time allotted at the scanning facility (for limitations on scanner availability, please see below). Overall, the scanning success rate in the present study was 98%, which is considerably higher than the 79% rate reported previously for infants < 3-months-of-age (Antonov et al., 2017). However, as described in detail below, data from twenty-four additional participants had to be excluded due to low image or segmentation quality, reducing the success rate to 54% (29/54).

Following these exclusions, twenty-nine infants (79 ± 10 days-of-age; F/M = 12/17) were included in the final analysis (please see Table 1 for demographic details). All participants were typical, healthy infants with no diagnosed incidence of neurological disease or disability. A clinical radiologist in Bangladesh and a pediatric neuroradiologist at Boston Children’s Hospital (BCH) examined the MRI images to ensure the absence of any malignant brain features. The study was approved by research and ethics review committees at BCH and The International Centre for Diarrhoeal Disease Research, Bangladesh (icddr,b).

**Table 1.** Subject Demographics.

|  |  |
| --- | --- |
| N | 29 |
| Sex (F/M) | 12/17 |
| Age (Days) | 79 ± 10 |
| Age Range (Days) | 65 - 98 |
| HAZ | -1.19 ± 1.0 |
| WAZ | -0.916 ± 1.0 |
| WHZ | 0.182 ± 0.91 |
| Monthly Income-to-Needs* | 5800 ± 3200 |
| Maternal Education (Years) | 6.45 ± 3.6 |

\*Monthly Income-to-Needs in Tk (Bangladeshi currency)/ number household members

### Challenges in Conducting an MRI Study in Dhaka, Bangladesh

Additional considerations for conducting the first MRI study in a particular low-resource setting (Dhaka, Bangladesh) are detailed in a previous publication (Turesky et al., 2019). The present study was a collaboration among many international facilities (BCH, Harvard University, University of Virginia, icddr,b, and the National Institute of Neuroscience and Hospital – NINSH). Main challenges included differences in languages (between English and Bengali) and time zones (11 hours between Boston and Bangladesh), which complicated coordination between sites. Additionally, political tensions at various times limited which personnel from the U.S. could safely travel to Bangladesh for in-person collaborations. Finally, because the scanner at NINSH is not connected to the internet, MRI data needed to be transferred across multiple sites before they were accessible to BCH.

Recruitment for the current study was conducted differently from studies in higher-resource settings. First, experienced senior investigators at icddr,b selected the Mirpur area of Dhaka from which to recruit participants. Mirpur was chosen for its population density (with over 1 million residents) and socioeconomic diversity. Within Mirpur, participants were recruited from three of the poorer neighborhoods (Kirkpatrick et al., 2015). Then, medical officers from icddr,b visited the homes of pregnant mothers to assess eligibility for and interest in enrollment. Eligible women who were willing to participate were escorted to a field site of icddr,b, established in Mirpur, for informed consent, and returned to this field site for behavioral testing. All eligibility and consent forms, study descriptions, questionnaires, and behavioral batteries were translated from English to Bengali for this study.

In preparation for infant data collection, NINSH staff undertook additional training, as prior to this study they had scanned infants only for clinical, but not for research, purposes. This distinction is important for a few reasons. For instance, sedation may be employed for clinical scans, but is usually not used for research scans in any setting because it poses an unnecessary risk of harm. Therefore, NINSH staff were taught techniques to facilitate MRI acquisition during natural sleep (Antonov et al., 2017; Dean et al., 2014; Hughes et al., 2017). To do so, staff first traveled to Boston to observe and assist in pediatric neuroimaging sessions. BCH staff subsequently visited Dhaka, where they collaborated with NINSH staff on a plan to prepare families for experimental sessions in such a way that maximized the likelihood that MR images could be collected while infants slept naturally. For example, infants were kept awake during the commute from their homes to the MRI facility to increase the likelihood that they would be ready to sleep when scan time started. Also, staff assisted caregivers during the process of putting the baby to sleep by minimizing surrounding noises, darkening the room that was utilized for this purpose, and minimizing immediate stressors for the participating families. This plan was critical because it addressed key challenges not often encountered in higher-resource settings. For instance, in higher-resource settings, infant scans may be scheduled around their naptimes to maximize the likelihood that they will sleep during the scan (Raschle et al., 2012). As infant naptimes are unstructured in Dhaka, staff did not have this advantage. This challenge was intensified by the limitations in scanner availability. Government-owned and used almost exclusively for clinical scans, the scanner was available for this study only between 7:00 am and 9:00 am. Importantly, many more challenges were encountered than were discussed here, but a previous report by our group details these (Turesky et al., 2019).

### Anthropomorphic Measures

Stunting, underweight, and wasting were assessed using height-for-age (HAZ), weight-for-age (WAZ), and weight-for-height (WHZ) scores, respectively. Trained, local staff measured height (in centimeters) and weight (in kilograms) at the time of the scan and these measures were converted into age- and sex-referenced and standardized z-scores (i.e., HAZ, WAZ, and WHZ), using the World Health Organization’s Anthro Plus software (version 3.2.2). These standardized z-scores represent growth curves that were constructed during the Multicentre Growth Reference Study, conducted between 1997 and 2003 by fitting longitudinal and cross-sectional measurements from 8440 infants (0-24 months) and children (18-71 months) from Brazil, Ghana, India, Norway, Oman and the U.S. using the Box-Cox-power-exponential method with cubic splines for curve smoothing. Importantly, the infants and children comprising this dataset were raised in healthy environments (including breastfeeding and absence of smoking) that are “likely to favour the achievement of their full *genetic* growth potential” (WHO Multicentre Growth Reference Study Group, 2006). Because of this, consistent deviations from these standard growth curves can be inferred to reflect environmental hazards during upbringing. Table 1 summarizes these measures in the final cohort.

### Measures of Poverty

Monthly family income, number of household members, and years of maternal education were reported at the time of enrollment in the study (i.e., shortly after birth or at age 2 months in the additional sample) through oral interviews with the mothers of the infants. The first two measures were used to compute the income-to-needs variable, which was used with years of maternal education as an estimate of poverty in later analyses. With an exchange rate of USD$1:Tk80 (Bangladeshi taka), the poorest family in the cohort earned Tk1300 or USD$16.25 per month per household member, while the wealthiest family in the cohort earned Tk13000 or USD$162.50 per month per household member. For context, the World Bank (https://data.worldbank.org/) set the international extreme poverty standard at roughly $57 per person per month (calculated from USD$1.9/day and assuming 30 days/month). Maternal education was measured as years of formal education, ranging from 0 to 10, with 0 indicating no formal education, 1-9 indicating number of grades passed, and 10 indicating education beyond the 9^th^ grade passed, in which degrees may be conferred.

### MRI Data Acquisition

All neuroimaging data were acquired while infants were sleeping naturally in a 3T Siemens MAGNETOM Verio scanner located at NINSH in Dhaka, Bangladesh (Fig. 2B). Structural T_1_-weighted magnetization-prepared rapid gradient-echo prospective motion-corrected (MPRAGE) scans were acquired with the following parameters: TR = 2520 ms, TE = 2.22 ms, 144 sagittal slices, 1 mm^3^ voxels, FOV = 192 mm. Resting-state functional MRI sequences were also acquired and are described in a previous report by our group (Turesky et al., 2019). T_2_-weighted structural images were also acquired, but data quality was substantially reduced due to motion, as this sequence was completed last and many infants awoke during this sequence. Therefore, most of these datasets were not usable for image processing purposes. Finally, diffusion-weighted imaging was performed and will be discussed in future reports.

### MRI Pre-Processing

Raw MPRAGE images (Fig. 3A) were submitted to the FreeSurfer recon-all pipeline for infant MRI images (Zöllei et al., 2017) for segmentation into cortical and subcortical gray and white matter regions and ventricles, all in native space. This pipeline (https://surfer.nmr.mgh.harvard.edu/fswiki/infantFS) takes a multi-atlas label fusion approach and selects relevant atlas information to use, depending on similarity to the brain to be segmented. The prior-based Bayesian method (Iglesias et al., 2012) relies on a training dataset of infant structural images, in which brain areas such as the amygdala were individually, manually segmented (de Macedo Rodrigues et al., 2015). Deformable Registration via Attribute Matching and Mutual-Saliency Weighting (DRAMMS) (Ou et al., 2010) is used to register atlas and subject data. It should be noted that there is still no consensus regarding the best segmentation pipeline for infant imaging (Makropoulos et al., 2018; Zhang et al., 2019). Segmentations were subsequently consolidated into binary gray matter (GM), white matter (WM), and cerebrospinal fluid (CSF) tissue maps for each participant (Fig. 3B) using in-house MATLAB 2015b (*MathWorks*) code.

**Figure 3.**
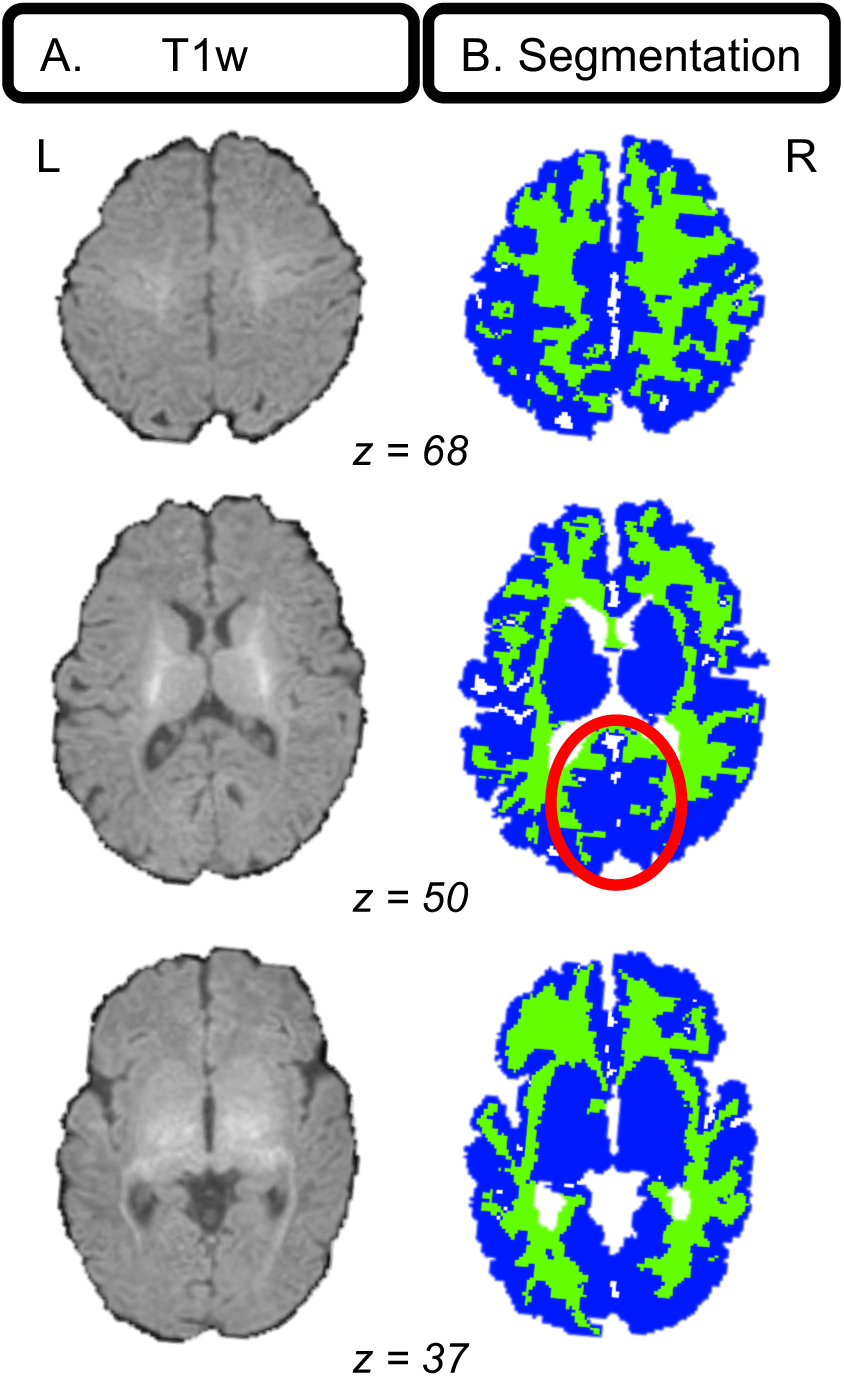
Example infant segmentation. (A) Three axial slices of a typical T_1_-weighted image from an infant whose segmentation had relatively accurate GM and WM labeling by a pediatric neuroradiologist. Notable are the low contrast delineating gray and white matter and thin cortex. (B) Segmentations for the T_1_-weighted image in A depicting gray (blue) and white (green) matter. Gray/white matter mislabeling is noticeable in medial occipital cortex (red circle).

### Segmentation Scoring

Segmentations were carefully evaluated by a pediatric neuroradiologist at BCH for accurate GM and WM labeling and rated according to a 0-10 scale with 0 = poor segmentation; 10 = excellent segmentation. Segmentations of adult and other infant brains acquired on MR scanners at BCH were used as baseline, independent examples of “poor” and “excellent” segmentations. Only segmentations greater than or equal to 7/10 were used in subsequent analyses. All segmentations were visualized using the Mango software package (http://rii.uthscsa.edu/mango/). After careful review, 24 of 53 participants were excluded due to poor image, leaving 29 infants with segmentation quality scores ≥7/10. Indeed, segmenting infant brains with high accuracy and precision is extremely challenging with currently available methods, given lower contrast-to-noise, higher rates of head motion, increased WM-CSF partial volume effects, and other obstacles compared with segmentations for other age groups (Makropoulos et al., 2018).

### Statistical Analyses

Total (absolute) intracranial G/WMV estimates were calculated from 29 participants whose segmentations were of relatively high quality by counting the number of voxels in each subjects’ G/WM masks; as the voxel size was 1 mm^3^, no additional calculation was needed to convert from units of voxels to mm^3^. Brain-anthropometry relationships were investigated by submitting GMV and WMV estimates to correlation analyses with HAZ, WAZ, and WHZ, while controlling for chronological age, sex (given sex differences found in earlier reports; e.g., (Choe et al., 2013)) and multiple comparisons (six tests; Bonferroni). In this semi-partial design, variance contributed by chronological age and sex was only removed from brain volume, but not anthropomorphic, measures because the latter were already standardized for age and sex. Resulting GMV and WMV residuals (i.e., adjusted G/WMV estimates) were subsequently tested for correlations with anthropometric indicators. To ensure that significant brain-anthropometry relationships were not confounded by head size or segmentation quality, head circumference and segmentation ratings provided by the pediatric neuroradiologist (please see above) were submitted to additional correlation analyses with anthropomorphic and adjusted G/WMV estimates; as with G/WMV estimates, head circumference was also adjusted for chronological age and sex prior to tests of correlations. Neither head circumference nor segmentation quality rating was correlated (p_unc_ > 0.05) with anthropomorphic or tissue volume estimates.

To test whether poverty is linked to anthropometry and brain structure, additional correlation analyses were performed between anthropomorphic and brain structural variables (adjusted for chronological age and sex) and income-to-needs and years of maternal education. Here, we only examined the anthropomorphic and brain structural variables that exhibited significant brain-anthropometry semi-partial correlations in the previous step and GMV due to previously documented income-to-needs and GMV associations (Betancourt et al., 2016; Hanson et al., 2013). To test whether poverty contributes unique variance to brain structure at two-three months-of-age, independent of the variance contributed by anthropomorphic estimates (or, more specifically, the underlying biological risk factors that shape these), variables from significant brain-anthropometry relationships were submitted to a hierarchical multiple linear regression with measures of poverty using SPSS 23. Measures of brain structure served as dependent variables, while anthropomorphic estimates and poverty measures served as independent variables; akin to the semi-partial correlations described earlier, these models controlled for chronological age and sex. Variance associated with chronological age and sex was removed in step 1; variance associated with anthropometry was estimated in step 2, and variance contributed by poverty was estimated in step 3. Separate regression models were tested for each combination of poverty measure (income-to-needs and years of maternal education) and significant brain-anthropometry pair variables revealed by the earlier semi-partial correlations.

## RESULTS

To examine the impact of biological risk factors on brain structure, we calculated anthropometry and estimated G/WMVs for each subject and tested semi-partial correlations with these variables. Next, measures of poverty, particularly income-to-needs and maternal education, were examined for relationships with anthropomorphic and brain structure measures using additional statistical testing.

### Anthropometry

Anthropomorphic estimates of stunting, underweight, and wasting were calculated; respectively, infants in the present cohort exhibited mean (± standard error) HAZ of −1.19 (±0.19), WAZ of - 0.916 (±0.19), and WHZ of 0.182 (±0.17). According to the World Health Organization (WHO) designated threshold of −2 for these three anthropomorphic estimates, eight infants exhibited stunting (2 females) and six infants exhibited underweight (1 female); no infants exhibited wasting.

### Brain-Anthropometry Relationships

Infants exhibited mean (± standard error) absolute GMV of 334392 (±5076) mm^3^ and WMV of 160806 (±2856) mm^3^. By comparison, Knickmeyer and colleagues (2008) estimated cortical and subcortical GMV at 261144 (±3220) mm^3^ (summed gray matter and subcortical estimate means and variances) and cortical WMV at 164,433 (±1901) mm^3^ in a U.S. sample of neonates (mean age: 0.75 ±0.31 months).

When examining relationships between anthropomorphic estimates and measures of brain structure, semi-partial correlations (adjusting brain volume estimates for chronological age and sex) were observed between HAZ and WMV (r = .52, p_corr_ < .05) and WAZ and WMV (r = .52, p_corr_ <.05). A semi-partial correlation between GMV and HAZ was also observed (r = .41; p_unc_ < .05); however, this relationship did not pass correction for multiple comparisons (six tests). WHZ was not correlated with GMV or WMV (Fig. 4).

**Figure 4.**
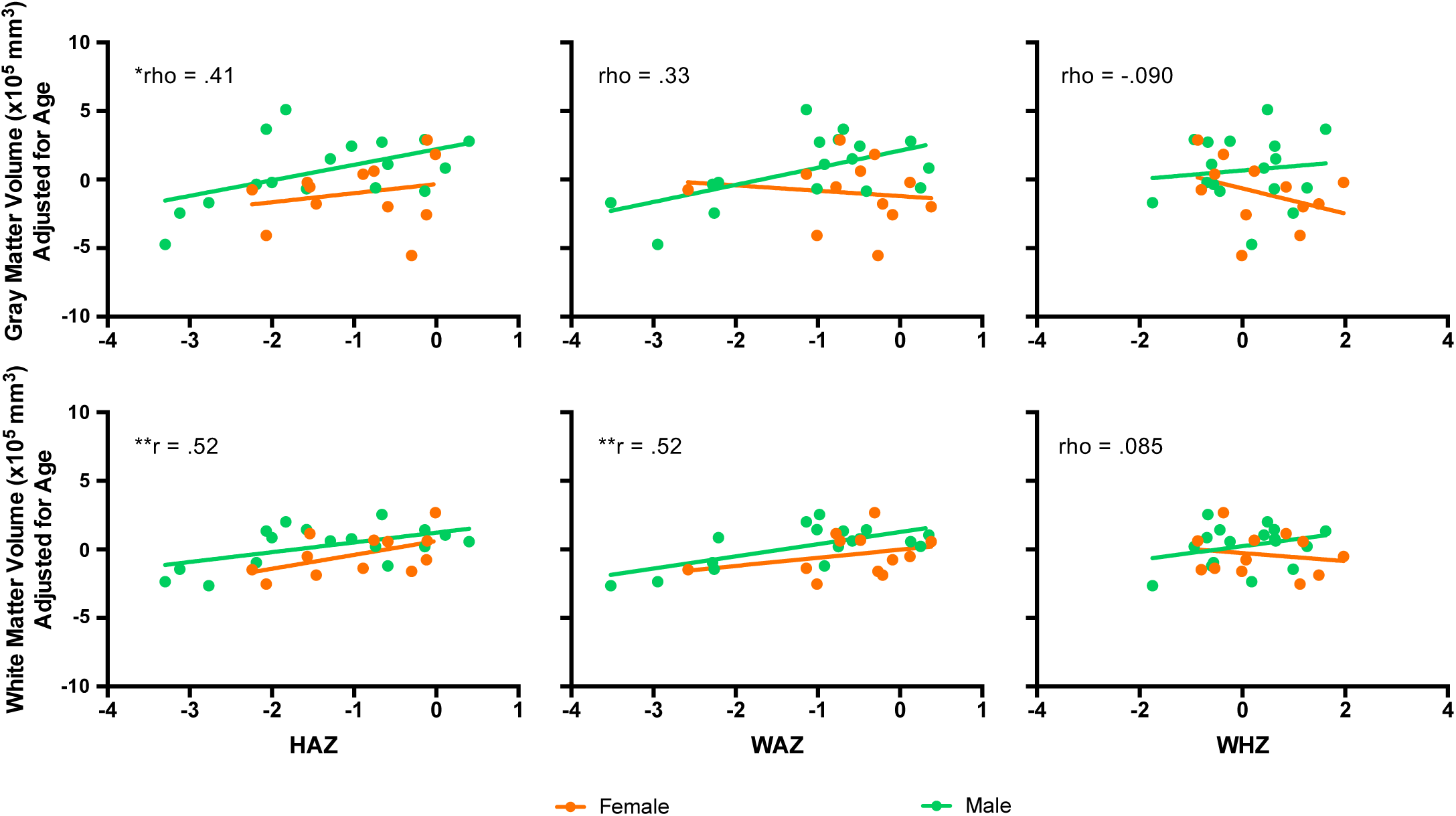
Brain-anthropometry relationships. Anthropometric indicators measured with HAZ, WAZ, and WHZ are depicted on x-axes and measures of gray matter volume (top row) and white matter volume (bottom row) are shown on y-axes for female (orange) and male (green) infants separately. Relationships were calculated using partial correlations, controlling for chronological age and sex. *p_unc_ < .05, **p_bonf_ < .05 (6 tests).

### Relationships between Measures of Anthropometry, Brain Volume, and Poverty

To examine whether measures of anthropometry and brain structure were associated with poverty, anthropomorphic estimates of HAZ and WAZ and chronological age- and sex-adjusted GMV and WMV estimates (i.e., the variables that survived correction in the previous analysis and GMV, as it has been associated with income-to-needs previously (Betancourt et al., 2016; Hanson et al., 2013)) were subsequently submitted to additional correlation analyses with two measures of poverty: income-to-needs and years of maternal education. No significant correlations were identified.

Hierarchical multiple linear regression was also performed, beginning with WMV as the dependent variable and chronological age and sex as the first independent variables (model 1: R^2^ =.080, F(2, 26) = 1.137, p = .336). In model 2, anthropomorphic variables of (A) HAZ (model 2A: R^2^ =.336, F(3, 25) = 4.214, p = .015) or (B) WAZ (model 2B: R^2^ = .348, F(3, 25) = 4.445, p = .012) were added. The addition of anthropomorphic estimates improved the model for HAZ (ΔR^2^ = .255, F(1, 25) = 9.615, p = .005) and WAZ (ΔR^2^ = .267, F(1, 25) = 10.251, p = .004).

Finally, model 3 incorporated (i) income-to-needs or (ii) years of maternal education into step 2 models for HAZ (model 3Ai: R^2^ = .447, F(4, 24) = 4.856, p = .005; model 3Aii: R^2^ = .359, F(4, 24) = 3.365, p = .025) and WAZ (model 3Bi: R^2^ = .395, F(4, 24) = 3.910, p = .014; model 3Bii: R^2^ = .348, F(4, 24) = 3.205, p = .030). The addition of income-to-needs improved only model 2a, which used HAZ (ΔR^2^ = .111, F(1, 24) = 4.839, p = .038), but this was not significant when controlling for the number of models generated. The alternative addition of years of maternal education did not improve the model for HAZ or WAZ.

## DISCUSSION

To examine the relationship between anthropometric indicators and brain structure, we conducted an MRI study in 2-3-month-old infants growing up in Dhaka, Bangladesh, an extremely low-resource setting where biological hazards are severe. Only a small sample, constituting 54 infants, was recruited because the challenges of conducting infant MRI in a low-resource area, at facilities that had not previously conducted infant MRI studies, needed to be realized and resolved before introducing a larger cohort. Despite the small sample size and other challenges (please see Methods), total gray and white matter volume (G/WMV) estimates were consistent with previous studies of infants scanned in a U.S.-based sample in the first few months of life (Gilmore et al., 2007; Knickmeyer et al., 2008). Examining the relationship between anthropometry and brain structure, WMV was positively correlated with height-for-age (HAZ) and weight-for-age (WAZ; while controlling for chronological age and sex). Thus, taller and heavier infants exhibited greater WMV. Effects were only observed for WMV; relationships between anthropometry and GMV were not significant after controlling for multiple comparisons. Also, weight-for-height (WHZ), the measure for wasting, was not associated with any measures of brain volume. In examining the role of poverty, no associations were observed between income-to-needs or maternal education and brain volumetric measures, suggesting that risk factors previously linked with poverty were not associated with brain volume pre- or peri-natally in this sample.

Key expectations for conducting infant MRI in Dhaka are detailed in a previous report (Turesky et al., 2019). Briefly, these challenges included lack of equipment and trained staff essential for undertaking infant MRI studies; limited scanner availability; unstructured infant naptime; the need for staff to recruit families in person due to limited communication channels; and coordination of investigators in Boston and research staff at two sites in Dhaka, which was further challenged by differences in time zone, differences in weekend and holiday schedules, and language barriers. Inferences drawn from these findings can also be problematic because infants in this low-resource setting are exposed to myriad risk factors, each of which may alter brain structure differently (e.g., Sheridan and McLaughlin 2014). This makes it difficult to match risk factors to their corresponding effects (a task that would be further complicated by interaction effects arising from multiple risk factors). Furthermore, interpretations of the findings presented here in the context of previous MRI studies of early life adversity must be done with the understanding that these previous studies were conducted in middle- and high-resource settings, without the multitude and severity of risk factors observed in Dhaka.

Results of the present study constituted correlations between HAZ, WAZ, and WMV after controlling for chronological age and sex, which supports the hypothesized link between anthropometry and brain structure. This link has a strong basis in cellular and molecular processes, in that many of the nutrients essential for neural growth are also essential for non-neural growth (e.g., manifesting in greater HAZ scores); e.g., zinc is involved in protein synthesis and nucleic acid metabolism, processes that occur in neural and non-neural domains. Conversely, lack of these nutrients (i.e., a form of malnutrition) would presumably derail widespread growth processes (Jensen et al., 2017). Despite this mechanism, the validity of anthropometric indicators in reflecting malnutrition in infants under six months-of-age has been questioned (Kerac et al., 2015), deepening the challenges associated with identifying malnutrition in this age group (Bhutta et al., 2017). It is important to note, however, that the results reported in the present study are correlational; causal mechanisms explaining the observed correlations were not directly tested.

The correlations between HAZ, WAZ, and WMV may alternatively be explained by underlying mechanism related to inflammation, which has been consistently linked to anthropometric indicators (Black et al., 2013; de Onis and Branca, 2016; Stewart et al., 2013) and white matter damage (Duggan et al., 2001; Hagberg et al., 2015; Hansen-Pupp et al., 2005). Especially germane to the present study, concentrations of pro-inflammatory markers measured in Bangladeshi infants during their first year of life are negatively associated with later neurodevelopmental outcomes (1-2-years-old) (Jiang et al., 2017, 2014), implying that inflammation may alter brain development in some way. Additionally, neurosonography studies in preterm infants partially corroborate the animal literature, in that mothers of infants with white matter injury had higher concentrations of inflammatory cytokines in prenatal amniotic fluid compared with mothers of infants without injury (Yoon et al., 1997). Further, repeated incidences of perinatal inflammation result in a greater likelihood of white matter injury, suggesting that the effects of inflammation on white matter appear to be cumulative (Korzeniewski et al., 2014). Although the presence or absence of white matter injury was inferred by evidence of associated pathologies (e.g., ventriculomegaly) or antecedents or counterparts of white matter injury (e.g., periventricular lesions), rather than directly measured, these results imply that white matter structure is vulnerable to inflammation in humans. Importantly, the observations made in humans have been corroborated and expounded by animal work. In mice, it was observed that systemic inflammation interfered with normative maturation of oligodendrocytes, and consequently central nervous system myelination, which manifested in reduced white matter microstructural connectivity and poorer performance on cognitive tasks (Favrais et al., 2011).

That effects in the present study were observed for white, but not gray, matter volumes parallels the predominance of studies reporting on the impact of inflammation on white matter, compared with the paucity on gray matter (Hagberg et al., 2015). Still, animal studies have also shown that malnutrition and inflammation alter cortical cell size, density of cortical dendritic spines, myelin production, number of synapses, and number of glial cells (Hagberg et al., 2015; Keunen et al., 2015; Levitsky and Strupp, 1995). While these cellular alterations likely alter measures of gray matter volume (e.g., Zatorre et al. 2012), it is possible that these effects are localized to specific brain regions and would not be detected when examining total volume (as done in the present study), or that altering total gray matter volume requires prolonged malnutrition or repeated inflammatory insults.

As part of the model proposed previously (Jensen et al., 2017) and adapted here (Fig. 1), we had also hypothesized a correlation between measures of poverty, such as income-to-needs or years of maternal education, and brain volumetric measures that would have been mediated by biological hazards (e.g., malnutrition or infection). However, as mediation requires an initial correlation between independent and depend variables, the absence of correlations between measures of poverty and GMV or WMV precluded a mediation analysis. To examine whether any of the observed variance in brain volume measures could be explained by poverty independently of the variance explained by chronological age, sex, and anthropomorphic measures, a hierarchical linear regression was performed. Neither the addition of income-to-needs nor maternal education improved the model explaining variance in brain structure (after controlling for multiple comparisons), suggesting that measures of poverty are not associated with brain structure by another, independent pathway (e.g., via psychosocial as opposed to biological risk factors), at least not at 2 months-of-age. These results diverge from previous studies of brain structure in infants, which have reported positive relationships between income-to-needs (or another income measure) and frontal and parietal lobe (Hanson et al., 2013), and cortical (Betancourt et al. 2016) and total gray matter volumes (Hanson et al., 2013). Importantly, though, the study by Hanson and colleagues (2013) was conducted on a cohort consisting of older infants (beginning at five months-of-age) and young children (up to 4 years-of-age), and the study by Betancourt and colleagues (2016), conducted on 1-month-old infants, restricted the sample to females. The sample size in the present study would have been too small to examine female infants alone.

There are three additional theoretical explanations for the null finding related to poverty. One could be that multiple insults from risk factors associated with poverty are required to effect detectable anatomical changes. That repeated inflammatory responses result in a greater likelihood of white matter injury supports this possibility (Korzeniewski et al., 2014). It is possible that sufficient insult has not accumulated to effect detectable anatomical changes in these two-month-old infants. A second possible explanation is that poverty-related alterations in brain structure are localized to specific brain regions and thus cannot be captured by examining measures of total tissue volume. A third possible explanation is that the anthropomorphic and brain volume variables examined may not be measured with sufficient accuracy and precision. Although only segmentations of relatively high quality were used, these segmentations were still of low quality relative to segmentations produced from structural images in adults and older children; this is perhaps not surprising as the cortex of an infant’s brain is thin and the contrast between gray and white matter is low. The difficulty of infant segmentation is well known (Makropoulos et al., 2016) and acknowledgment of this has led to nonconformity in infant segmentation pipelines (Makropoulos et al., 2018; Zhang et al., 2019). With regard to anthropometry, infant weight, for instance, can vary considerably depending on quantities eaten and recency of quantities expelled. Lack of reproducibility in anthropometric indicators for infants under six months-of-age has led to doubt over the reliability of these measures (Kerac et al., 2015).

Intriguingly, the relationship between anthropometry and poverty is age-dependent (Victora et al., 2010), with children from low-resource countries exhibiting lower anthropometric indicator scores on average as they grow up. If trajectories of brain structure parallel anthropometric indicator trajectories, as white matter volume does according to the present study, then poverty-related brain volume differences would be smaller at lower ages. This may be due to a few insults or protective factors such as nutrition from breastmilk (Bhutta et al., 2017). Most of the infants in the present sample were breastfed.

In sum, this pilot study demonstrated that it is feasible to conduct an infant MRI study of brain structure in a low-resource setting, such as Dhaka, Bangladesh. Anthropometric indicators, which have been associated with poor neurocognitive outcomes, correlated with measures of WMV. However, measures of poverty tested here (income-to-needs and maternal education) were not related to anthropometric indicators or measures of G/WMV in infants at two months-of-age. Although these results should be viewed as preliminary, given the small sample size and relatively low image segmentation quality, they provide the first evidence of a link between stunting, underweight, and early brain structure.

## Acknowledgments

Funding for the study was provided by research grants from the Bill & Melinda Gates Foundation to CA Nelson [OPP1111625] and WA Petri [OPP1017093], and research grants to WA Petri from the Henske Foundation and the NIAID (R01 AI043596-17). We thank the families who participated in the study, the staff at icddr,b who facilitated data collection, and Uma Nayak for managing the database.

